# Targeting intracellular Neu1 for Coronavirus Infection Treatment

**DOI:** 10.1101/2022.09.09.507342

**Authors:** Darong Yang, Yin Wu, Isaac Turan, Joseph Keil, Kui Li, Michael H Chen, Runhua Liu, Lizhong Wang, Xue-Long Sun, Guo-Yun Chen

## Abstract

There are no effective therapies for COVID-19 or antivirals against SARS-CoV-2. Furthermore, current vaccines appear less efficacious for new SARS-CoV-2 variants. Thus, there is an urgent need to better understand the virulence mechanisms of SARS-CoV-2 and the host response to develop therapeutic agents. Here, we show host Neu1 regulates coronavirus replication by controlling sialylation on coronavirus nucleocapsid protein. Coronavirus nucleocapsid proteins in COVID-19 patients and in coronavirus HCoV-OC43-infected cells were heavily sialylated; this sialylation controlled the RNA binding activity and replication of coronavirus. Neu1 overexpression increased HCoV-OC43 replication, whereas Neu1 knockdown reduced HCoV-OC43 replication. Moreover, a newly developed Neu1 inhibitor, Neu5Ac2en-OAcOMe, selectively targeted intracellular sialidase, which dramatically reduced HCoV-OC43 and SARS-CoV-2 replication in vitro and rescued mice from HCoV-OC43 infection-induced death. Our findings suggest that Neu1 inhibitors could be used to limit SARS-CoV-2 replication in patients with COVID-19, making Neu1 a potential therapeutic target for COVID-19 and future coronavirus pandemics.

## Introduction

The ongoing pandemic coronavirus disease 2019 (COVID-19) caused by the severe acute respiratory syndrome coronavirus 2 (SARS-CoV-2) currently affects millions of lives worldwide, posing an overwhelming burden on global health systems and economies. Although vaccines are available, they appear to be less efficacious against newly emerging variants of the virus (Abdool Karim and de Oliveira, 2021; Chen et al., 2021; Grubaugh et al., 2021; Madhi et al., 2021; Priesemann et al., 2021; Schultze and Aschenbrenner, 2021), and SARS-CoV-2 continues to ravage many communities worldwide. Furthermore, SARS-CoV-2 is likely to become endemic (Phillips, 2021), leading to the emergence of vaccine-resistant variants and reinforcing the need for development of antiviral therapeutic agents. SARS-CoV-2 is a β-coronavirus with a large (30 Kb) positive strand RNA genome. There currently exist no antiviral drugs that specifically target SARS-CoV-2, similar to the situation that occurred during the 2002 SARS outbreak and in many other human coronaviruses infections. A combination of hydroxychloroquine and azithromycin might be effective (Gautret et al., 2020), although evidence on the safety and efficacy of these therapies is limited (Cavalcanti et al., 2020). Additionally, remdesivir may modestly accelerate recovery time (Wang et al., 2020), and has been approved by the FDA for treatment of hospitalized COVID-19 patients over the age of 12. Many of the monoclonal antibody treatments that effective with past variants do not demonstrate efficacy with the Omicron variant and evidence on the safety and efficacy of the novel COVID-19 oral antiviral Molnupiravir is also limited (Fischer et al., 2021; Gordon et al., 2021; Imran et al., 2021; Kabinger et al., 2021; Malone and Campbell, 2021; Painter et al., 2021). More recently, Pfizer’s antiviral Paxlovid (nirmatrelvir and ritonavir) was just approved by the FDA for use in children 12 and above, however, the efficacy against SARS-CoV-2 variants is still unclear. Thus, the development of novel therapeutics against SARS-CoV-2 remains a top priority for combating the current pandemic and future coronavirus outbreaks.

The coronavirus genome encodes four major structural proteins: the spike (S), nucleocapsid (N), membrane (M), and envelope (E) proteins, all of which are required to produce a structurally complete viral particle (Gussow et al., 2020; Ye et al., 2020). While our knowledge of SARS-CoV-2 pathogenesis is expanding rapidly, most studies have only focused on a few proteins, such as S protein, and their interaction with the host immune system. Less is known about mechanisms that globally control host and viral RNA synthesis during infection. N protein is one of the most abundant structural proteins, plays key roles in the regulation of viral replication and virion assembly, and is a major immunogen in coronavirus infection-induced disease. Therefore, N protein is an attractive target for diagnosis and treatment strategies against coronaviruses including SARS-CoV-2 (Gussow *et al*., 2020; Lang et al., 2020). Overall, the mechanisms by which specific cellular genes are induced and regulated during SARS-CoV-2 infection are poorly understood. Knowledge of how host defense genes are controlled might be key to understanding COVID-19 disease pathogenesis.

Sialylation, the most frequent modification of proteins and lipids, is the addition of sialic acids (a family of nine-carbon acidic monosaccharides) to the terminal residues of glycoproteins and glycolipids. This modification is important in self-nonself discrimination(Chen et al., 2009), phagocytosis of cancer cells by macrophages(Barkal et al., 2019), determination of immunoglobulin E allergic pathogenicity(Shade et al., 2020) and bacterial infection(Chen et al., 2014b; Wu et al., 2020). The sialylation level of a cell is largely dependent on the activity of two kinds of enzymes: sialyltransferases that add sialic acid residues to glycolipids or glycoproteins and sialidases that remove sialic acid residues from glycolipids or glycoproteins (Chen *et al*., 2014b). Sialidase inhibitors are useful tools for studying sialidase function and serve as drugs for sialidase-related diseases, such as viral infection. Inhibition of viral neuraminidase activity has been successfully utilized as a therapeutic approach for influenza infection (Glanz et al., 2018). Tamiflu (oseltamivir) and Relenza (Zanamivir), which are approved for treatment of influenza A and B, have almost no effect on human sialidases (neuraminidases) but are potent inhibitors of neuraminidase activity of the influenza NA protein (Glanz *et al*., 2018; Hata et al., 2008). Here, we investigated whether N protein is sialylated and whether sialylation influences the biological activity of N protein and developed novel sialidase inhibitors for treatment of coronavirus infection.

## Results

### Sialylation on coronavirus N protein

Glycans on virus play an important role in virus infection, immune evasion and immune modulation (Vigerust and Shepherd, 2007). Coronavirus S, E and M proteins are glycosylated, although N- and O-link glycosylation were found on the N protein in vitro overexpress system (Supekar et al., 2021), it is not clear whether the N protein from virion is glycosylated or not (Shajahan et al., 2021). Human coronavirus OC43 (HCoV-OC43) is a β-coronavirus responsible for mostly mild respiratory symptoms, thus, we analyzed the N- and O-glycans on N protein from HCoV-OC43 virion by mass spectrometry, N protein from HCoV-OC43 virion is also heavily glycosylated, containing both N- and O-linked glycans (Figure. S1). COVID-19 is caused by coronavirus SARS-CoV-2(Hu et al., 2021). To determine whether sialylation occurred on N protein from both coronavirus HCoV-OC43 and SARS-CoV-2, we performed a lectin blot with samples immunoprecipitated from the serum of COVID-19 patients and healthy controls and cell lysates infected with HCoV-OC43 with anti-N protein antibodies. The immunoprecipitated samples treated with or without sialidase and then separated by SDS-PAGE. As shown in Fig. 1a, b, N protein from both patients with COVID-19 and HCoV-OC43-infected cells was heavily sialylated. Sialic acid was mostly attached in α2, 6 linkage on N protein (Fig. 1a, b), and N protein sialylation was confirmed by sialidase treatment (Fig. 1b). Sialylation was also observed on SARS-CoV-2-N protein expressed in HEK293T cells (Fig. 1c), HCoV-OC43-N protein in THP-1 cells (Fig. 1d) and HCoV-OC43 virion (Fig. 1e).

**Figure 1.**
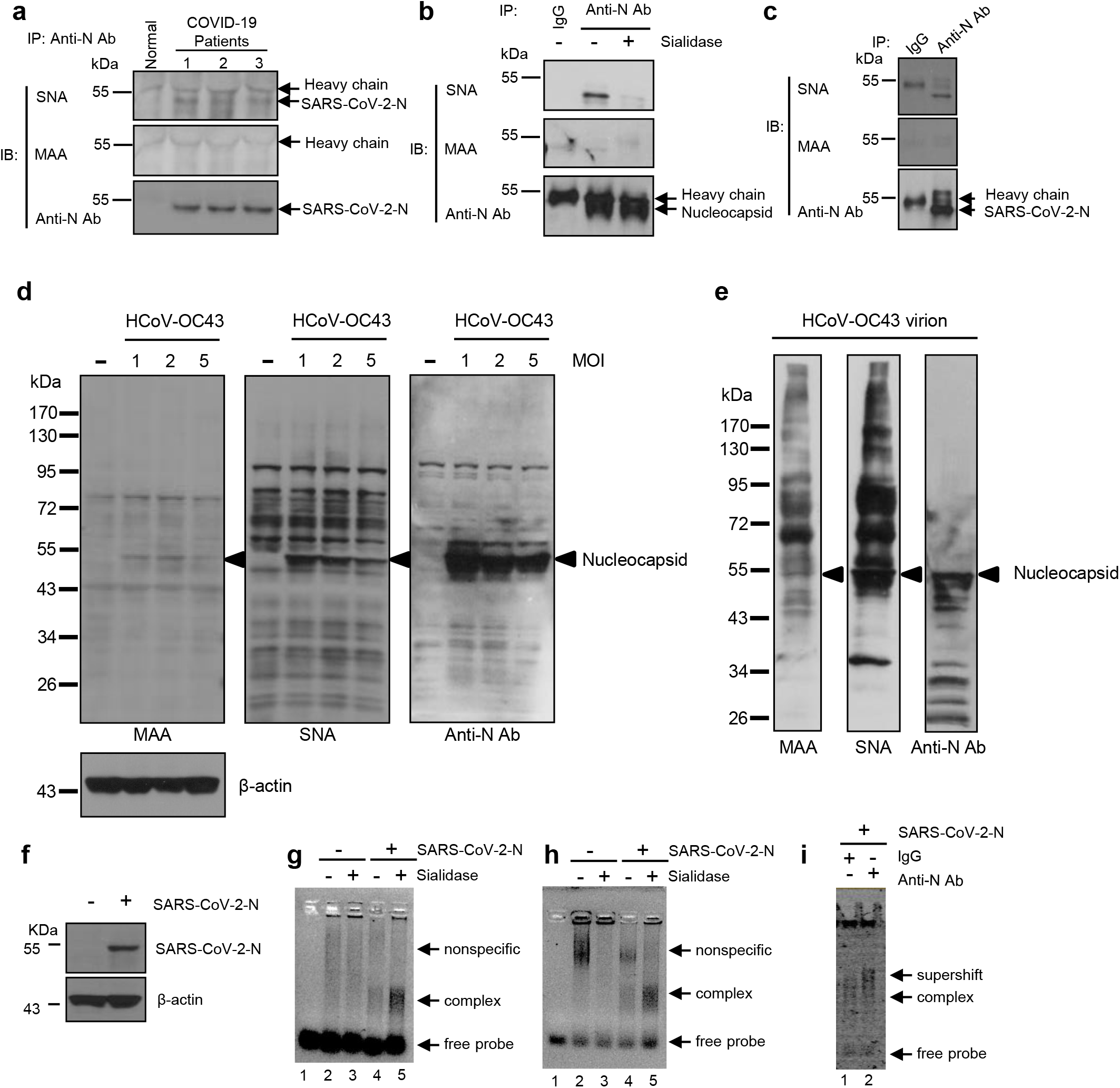
Sialylation on coronavirus nucleocapsid (N) protein is critical for its RNA binding activity. **a**–**c**, Immunoprecipitated concentration of nucleocapsid protein using the sera of COVID19 patients (**a**), cell lysates from HCoV-OC43-infected THP-1 cells (**b**), or HEK293T cell lysates overexpressing SARS-CoV-2 nucleocapsid (**c**), followed by immunoblot analysis of sialylation using biotin-MAA (α2,3-linkage), biotin-SNA (α2,6-linkage) lectins, biotin-anti-SARS-CoV-2-N (**a**), anti-HCoV-OC43-N or anti-SARS-CoV-2-N antibodies (**c**). **d**, Viral nucleocapsid is sialylated in host cell. Immunoblot analysis of proteins in THP-1 cells using biotin-MAA and biotin-SNA lectins and anti-HCoV-OC43-N (nucleocapsid) antibody. β-actin was used as the internal control. Nucleocapsid indicated with arrowhead. **e**, Viral nucleocapsid is sialylated in HCoV-OC43 virion. Immunoblot analysis of the sialylation of nucleocapsid (indicated with arrowhead) in HCoV-OC43 virion. **f**, Immunoblot analysis of the SARS-CoV-2 nucleocapsid protein in HEK293T cells. Cells were transfected with a construct expressing SARS-CoV-2 nucleocapsid (SARS-CoV-2-N) for 48 hours, and cell lysates were detected with anti-SARS-CoV-2-N and β-actin antibodies. **g**, **h**, Gel mobility shift assay of the 32-mer ssRNA (**g**) or 32-mer ssDNA (**h**). The probe was incubated with no cell lysates (lane 1) or lysates with the treatment indicated (lanes 2-5). Data are representative of three independent experiments. **i**, antibodies shift assay, 32-mer ssDNA was incubated with cell lysates after transfection with SARS-CoV-2 N protein expression vector and then added IgG (lane 1) or anti-N protein antibodies (lane 2) and separated in a 7% acrylamide gel. Data are representative of three experiments.

### The role of sialylation in coronavirus N protein

Since the primary role of N protein is to assemble with genomic RNA into the viral RNA-protein complex(Mu et al., 2020), we investigated whether the sialylation on N protein affects its RNA binding activity. To assess the nucleic acid-binding affinity of N protein, we conducted nucleic acid-binding assays in the presence of a 32-mer stem-loop II (32m) motif single-stranded RNA (ssRNA) and its 32-mer ssDNA mimic. The 32m ssRNA is a highly conserved sequence among coronaviruses and has been used to map the putative RNA-binding domain of SARS-CoV N protein(Huang et al., 2004; Robertson et al., 2005). For nucleic acid-binding assays, we lysed HEK293T cells 48 hours after transfection with SARS-CoV-2-N protein expression vector (Fig. 1f) and then treated the cells with or without sialidase. SARS-CoV-2-N protein formed a strong complex with 32-mer ssRNA (Fig. 1g) and 32-mer ssDNA (Fig. 1h), which was supershifted in the presence of anti-N protein antibodies (Fig. 1i), indicating that this complex is specific for N protein. As expected, HEK293T cell lysates transfected with empty vector did not form a complex with 32-mer ssRNA and ssDNA (Fig. 1g, h). Furthermore, 32-mer ssDNA and ssRNA bound to sialidase-treated N protein dramatically increased (Fig. 1g, h). These findings indicate a significant increase in N protein RNA binding activity after sialidase treatment, supporting the critical role of N protein sialylation in RNA binding.

### A critical role of Neu1 in coronavirus replication

The sialylation level of a cell is largely dependent on the activity of two kinds of enzymes: sialyltransferases are responsible for adding sialic acid residues to glycolipids or glycoproteins, while sialidases are responsible for removing sialic acid residues from glycolipids or glycoproteins(Chen *et al*., 2014b). We evaluated the contribution of endogenous sialidases to the sialylation of N protein using THP-1 cell lines. Real-time PCR (Fig. 2a) and western blot analysis (Fig. 2b) indicated the expression of Neu1 but not Neu2, Neu3 or Neu4 was significantly increased after infection with HCoV-OC43 for 72 hours. Notably, NEU1 is also upregulated in COVID-19 patients(Formiga et al., 2020). In addition, N protein associated with Neu1 in HCoV-OC43-infected cells (Fig. 2c).

**Figure 2.**
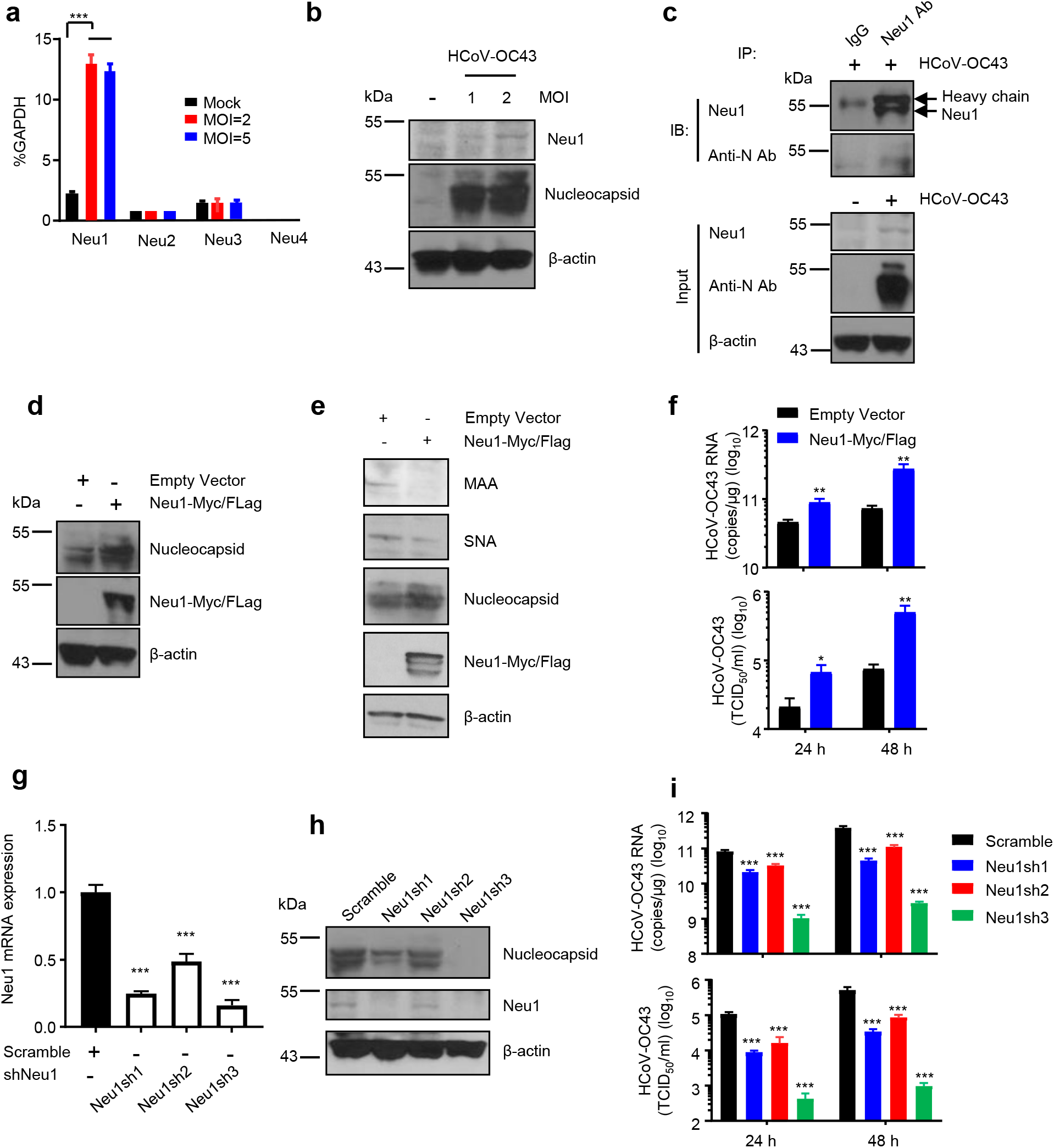
A critical role of Neu1 in coronavirus replication. **a**, The mRNA levels of Neu1, Neu2, Neu3 and Neu4 normalized by GAPDH in THP-1 cells with or without HCoV-OC43 infection for 72 hours. **b**, Immunoblot analysis of Neu1 in naïve and HCoV-OC43 infected THP-1 cells. β-actin was used as the internal control. **c**, N protein associated with endogenous Neu1 in HCoV-OC43-infected THP-1 cells (48 hours post-infection). **d**, Immunoblot analysis of Neu1 and viral nucleocapsid in Neu1 overexpressing THP-1 cells. **e**, Overexpression of Neu1 dramatically reduced the levels of sialylation on viral nucleocapsid in HCoV-OC43-infected 293T cells. 293T cells were infected with HCoV-OC43 (MOI = 1) for 24 hours, followed by transfection of empty vector or the construct expressing Neu1-Myc/Flag for 24 hours. Immunoblot analysis of the indicated targets. **f**, The levels of intracellular viral RNA (upper) and extracellular viral titers (lower) in control and Neu1 overexpressing THP-1 cells. **g**, The efficiency of Neu1 knockdown in THP-1 cells. RT-qPCR analysis of Neu1 mRNA abundance in THP-1 cells transduced with either scrambled shRNA or shRNAs targeting Neu1. **h**, Immunoblot analysis of Neu1 and viral nucleocapsid in scrambled and Neu1 knockdown THP-1 cells. **i**, The levels of intracellular viral RNA (upper) and extracellular viral titers (lower) in scrambled and Neu1 knockdown THP-1 cells. Data are representative of three (**a**-**c**) or two (**d**-**i**) independent experiments. Data are shown as mean ± SD. *p < 0.05; **p < 0.01; ***p < 0.001. Analysis was performed using two-way ANOVA.

Coronavirus N protein is a multifunctional RNA-binding protein necessary for viral replication(Mu *et al*., 2020). Since we determined the sialylation on N protein affects its RNA binding activity, we investigated whether this sialylation affects virus replication. Viral infection was quantified by real-time quantitative PCR (RT-qPCR) with primers targeting the coding region of the viral N gene. RNA was collected from cells at indicated time points after viral challenge, and viral transcripts were quantified. Supernatants were also processed for quantification of viral titer by 50% tissue culture infective dose (TCID_50_) assay(Liu et al., 2014). We made THP-1 stable cell lines overexpressing Neu1 (Fig. 2d), the sialylation on N protein was significantly decreased in the cells overexpressing Neu1 compared with empty vector control cells (Fig. 2e). The replication of HCoV-OC43 was more than 10-fold higher at the level of viral transcripts and viral titers in cell culture supernatants (Fig. 2f) of cells overexpressing Neu1 than in cells expressing empty vector 48 hours after viral challenge. By contrast, the replication of HCoV-OC43 was more than 100-fold lower at the level of viral transcripts and viral titers in the cell culture supernatants (Fig. 2i) of cells overexpressing shRNA for Neu1 (Fig. 2g, h) than in cells expressing scrambled shRNA. Compared with scrambled shRNA, Neu1sh3 significantly decreased HCoV-OC43 replication more than Neu1sh1 and Neu1sh2, consistent with the knockdown efficiency of shRNA (Fig. 2g, h and i). N protein levels were also significantly decreased in Neu1 knockdown cells (Fig. 2h) but dramatically increased in Neu1 overexpressing cells (Fig. 2d). These data indicate host Neu1 is a regulator of HCoV-OC43 replication in THP-1 cells.

### Sialidase inhibitors library screening

The significantly reduced HCoV-OC43 replication in Neu1 knockdown THP-1 cells suggests endogenous sialidase plays a key role in HCoV-OC43 replication and thus may be a valuable therapeutic target. To test this concept, we set up a sialidase inhibitor library which included FDA approved influenza sialidase inhibitors: Zanamivir and oseltamivir; cell surface sialidase inhibitors: Neu5Gc2en and Neu5Ac2en9N3; and newly synthesized cytoplasm sialidase inhibitors which are expected to have higher cell membrane permeability and higher cellular uptake than the parent compound, the active form of the inhibitor is subsequently released via hydrolysis by esterases (Schönemann et al., 2016) in cytosol: Neu5Ac2en-OMe, Neu5Ac2en-OAcOMe, and Neu5Ac2en9N3-OAcOMe. We first assessed these sialidase inhibitors for antiviral activity against HCoV-OC43 in vitro. THP-1 cells treated with these inhibitors were challenged with HCoV-OC43 for 2 hours, RNA was collected from cells, and viral transcripts were quantified 72 hours after viral challenge (Fig. 3a, b upper). Supernatants were also processed at 72 hours for quantification of viral titer by TCID_50_ assay (Fig. 3a, b lower). Three of the tested sialidase inhibitors significantly repressed viral replication (Fig. 3a, b). Among them, Neu5Ac2en-OAcOMe showed the highest antiviral activities (Fig. 3a), which were dose-dependent.

**Figure 3.**
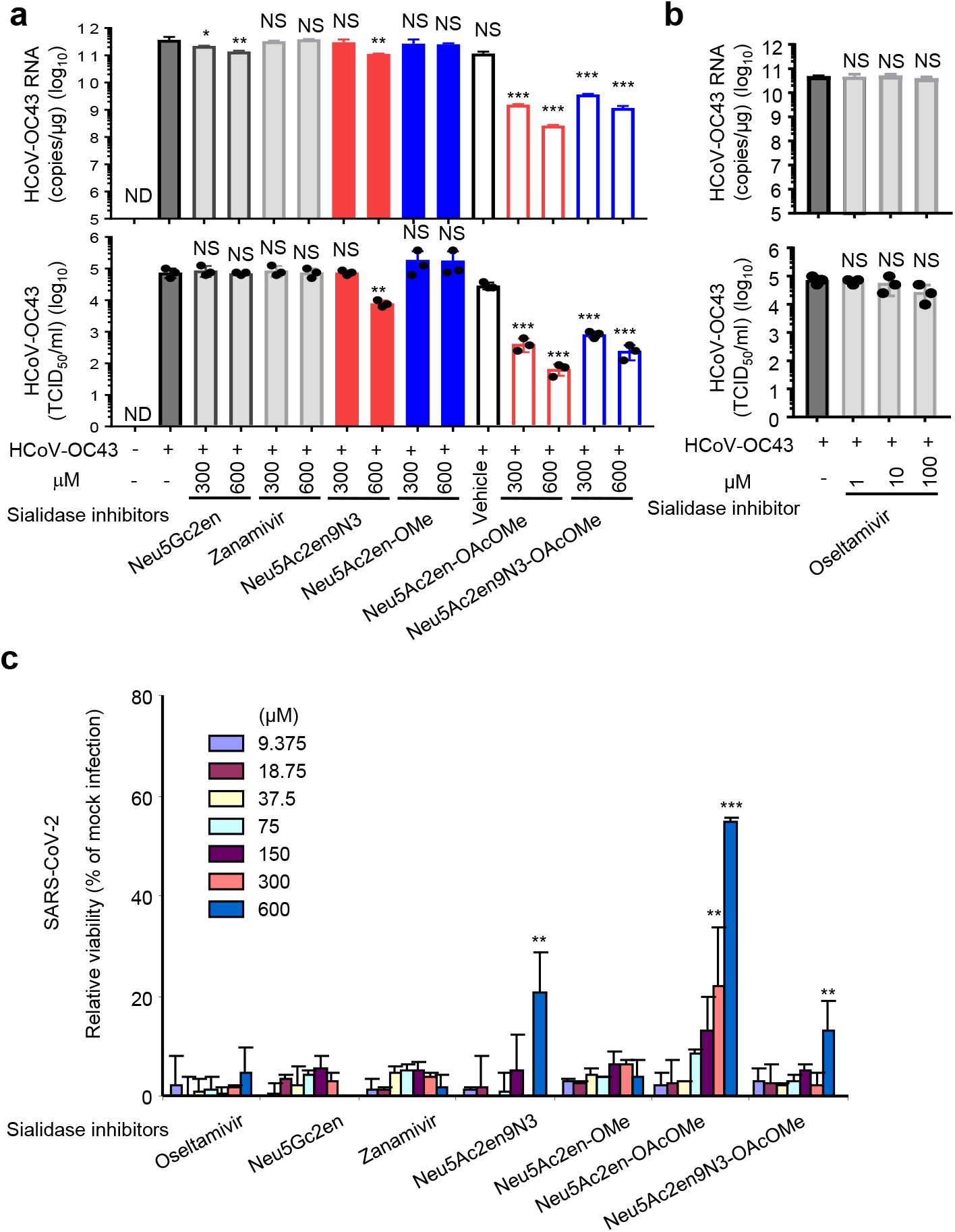
Screen for sialidase inhibitors suppressing coronavirus propagation. **a**, **b**, The intracellular viral RNA levels (upper) and viral titers (lower) in the supernatant of HCoV-OC43 (MOI = 2) infected THP-1 cells treated with sialidase inhibitors Neu5Gc2en, Zanamivir, Neu5Ac2en9N3, Neu5Ac2en-OMe, Neu5Ac2en-OAcOMe, Neu5Ac2en9N3-OAcOMe (**a**) or oseltamivir (**b**). **c**, Vero 76 cells were treated with the sialidase inhibitors and infected with SARS-CoV-2 for 48 hours; then cell viability was measured. Data are shown as mean ± SD and are representative of three (**a, b**) independent experiments or the two replicate screens (**c**). *p < 0.05; **p < 0.01; ***p < 0.001. NS, not significant. ND, not detected. Analysis was performed using one-way ANOVA.

### Screening for sialidase inhibitors that block SARS-CoV-2 replication

We next used a high-throughput screen (HTS) assay to test whether these inhibitors have similar effects on the replication of SARS-CoV-2 and used cell viability 48 hours post-SARS-CoV-2 infection as a readout. Neu5Ac2en-OAcOMe showed the best protection (Fig. 3c), while viral neuraminidase inhibitors Zanamivir and oseltamivir did not show inhibitory activity (Fig 3a, b and c). These findings are in agreement with previous reports that Zanamivir and oseltamivir have limited potency against all human sialidases(Hata *et al*., 2008).

### Effects of Neu5Ac2en-OAcOMe on coronavirus life cycle

To further evaluate the antiviral activity of sialidase inhibitors, we chose Neu5Ac2en-OAcOMe which showed the highest antiviral activities and first evaluated whether Neu5Ac2en-OAcOMe acts on the virus binding and entry steps of the viral life cycle. We treated THP-1 cells with Neu5Ac2en-OAcOMe for 2 hours and then infected the cells with HCoV-OC43 at 4°C or 37°C. Cells incubated at 4°C were collected 1 hour post-infection and cells incubated at 37°C were collected 2 hours post-infection. Intracellular viral RNA was quantified by RT-qPCR. As shown in Fig. 4a, viral loads were similar among cells treated with different amounts of Neu5Ac2en-OAcOMe, indicating that Neu5Ac2en-OAcOMe treatment did not affect virus binding and entry steps of the viral life cycle.

**Figure 4.**
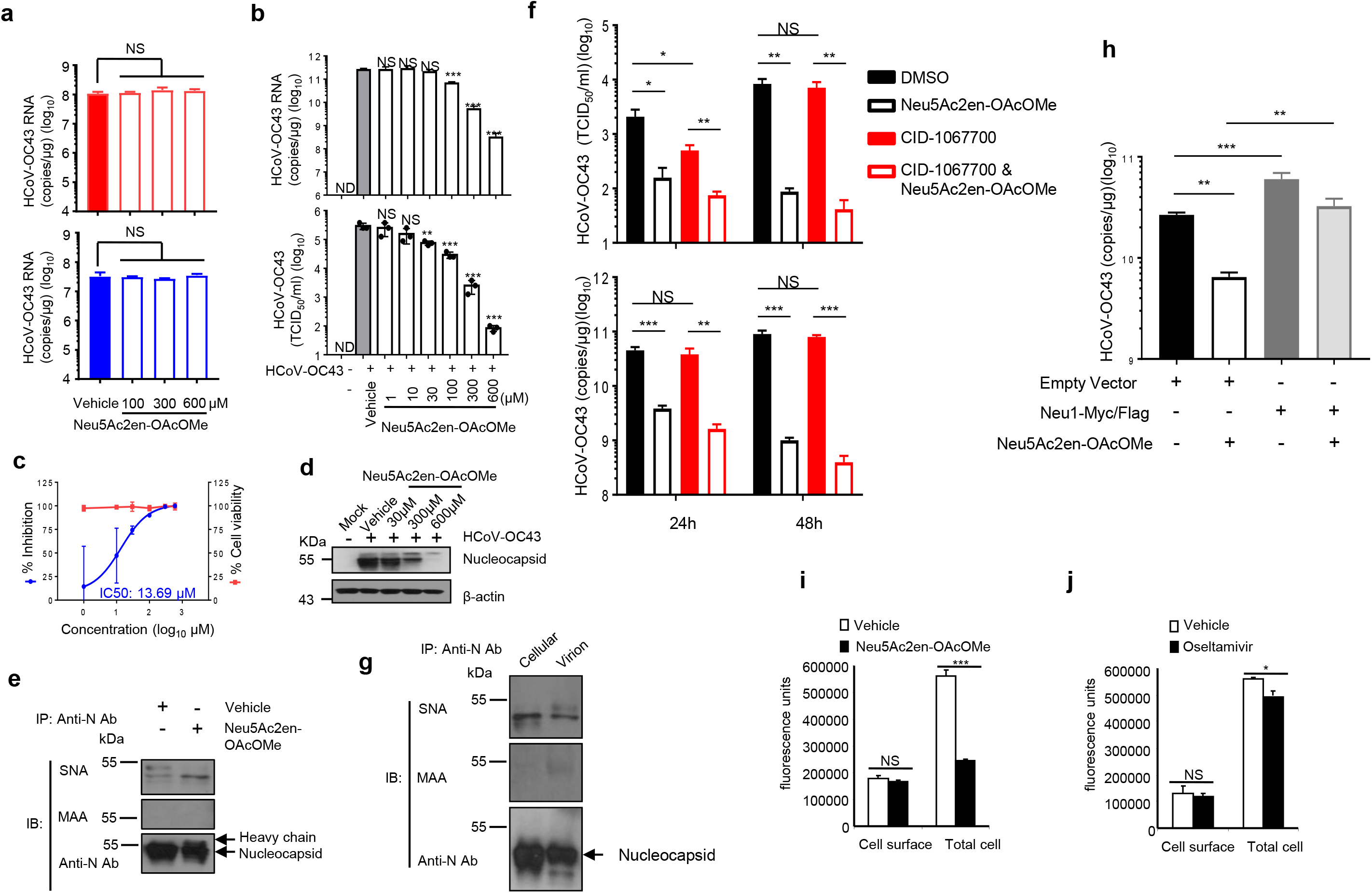
Neu5Ac2en-OAcOMe exerts anti-viral effects in human cell models. **a**, THP-1 cells were pretreated with vehicle or Neu5Ac2en-OAcOMe and were inoculated with HCoV-OC43 (MOI = 2) at either 4°C (upper) or 37°C (lower). Intracellular viral RNA was analyzed by RT-qPCR. **b**-**d**, HCoV-OC43 (MOI = 2) infected THP-1 cells were treated with a gradient concentration of Neu5Ac2en-OAcOMe for 72 hours. Intracellular viral RNA (**b**, upper), extracellular progeny virus yields (**b**, lower and **c**) and intracellular nucleocapsid (**d**) were analyzed by RT-qPCR, TCID_50_ assay and immunoblot, respectively. The inhibitory and cytotoxic curves (**c**) were obtained using the data from the lower panel in **b** and cell viability measured by MTS assay. CellTiter 96® AQueous One Solution Cell Proliferation Assay (MTS) (Promega, G3582) was performed according to the kit manual. **e**, HCoV-OC43-infected THP-1 cells were treated with vehicle or Neu5Ac2en-OAcOMe for 24 hours. Immunoprecipitation was performed to capture nucleocapsid using anti-HCoV-OC43-N antibody. Sialylation was detected with biotin-MAA and biotin-SNA lectins. **f**, CID-1067700 did not reverse the decrease in intracellular viral RNA and extracellular viral production triggered by Neu5Ac2en-OAcOMe. THP-1 (MOI = 1) cells were inoculated with HCoV-OC43 virus for 2 hours, and then CID-1067700 and Neu5Ac2en-OAcOMe were added after removal of inoculum. The levels of intracellular viral RNA (upper) and virus release (lower) were analyzed by RT-qPCR and TCID_50_ assay, respectively (72 hours post-infection. **h**, Neu1 overexpression reversed the inhibition of Neu5Ac2en-OAcOMe on HCoV-OC43 infection. HeLa cells transfected with empty vector or Neu1 expression vector were inoculated with HCoV-OC43 virus (MOI = 1), and then treated with or without Neu5Ac2en-OAcOMe for 48 hours. The levels of intracellular viral RNA were analyzed by RT-qPCR. **i**, **j**, Sialidase activity after incubation with Neu5Ac2en-OAcOMe (**i**) or oseltamivir (**j**). Data are representative of at least three (**a**-**e**, **g**) or two (**f**, **h**-**j**) independent experiments. *p < 0.05; **p < 0.01; ***p < 0.001. NS, not significant. ND, not detected. Analysis was performed using one-way ANOVA (**a**-**b**) or unpaired Student’s t-test (**f**, **h**-**j**).

To determine whether Neu5Ac2en-OAcOMe affects post-entry steps of the viral life cycle, we infected THP-1 cells with HCoV-OC43 for 2 hours and then treated the cells with Neu5Ac2en-OAcOMe. We quantified intracellular viral RNA and viral titers in the cell culture supernatants 72 hours post-infection. Neu5Ac2en-OAcOMe treatment significantly decreased viral replication in THP-1 cells at the level of viral transcripts (Fig. 4b, upper) and viral titers in cell culture supernatants (Fig. 4b, lower) in a dose-dependent manner, with an IC_50_ of 13.69 μM (Fig. 4c). N protein levels were also significantly decreased in Neu5Ac2en-OAcOMe-treated cells (Fig. 4d). Compared with vehicle-treated cells, Neu5Ac2en-OAcOMe-treated cells showed increased sialylation of HCoV-OC43 N protein (Fig. 4e). Importantly, Neu5Ac2en-OAcOMe was non-toxic at all tested concentrations in MTS assay (Fig. 4c) and did not induce cytokine production in THP-1 cells (Figure. S2). Moreover, CID-1067700, a competitive inhibitor of Rab7 activation that can block β-coronavirus egress(Ghosh et al., 2020), moderately inhibited HCoV-OC43 release, not viral RNA replication, at 24 hours post-infection. CID-1067700 also did not reverse the decrease in intracellular viral RNA and extracellular viral production triggered by Neu5Ac2en-OAcOMe treatment (Fig. 4f). In addition, similar levels of sialylation were observed on cellular N protein and N protein in HCoV-OC43 virion (Fig. 4g), indicating that virus budding did not affect sialylation of N protein. Collectively, these results demonstrated that viral RNA synthesis is a target of Neu5Ac2en-OAcOMe.

To rule out the possibility that Neu5Ac2en-OAcOMe induced protein degradation, we treated the cells with MG132. As shown in Figure. S3a, inhibiting proteasome activity with MG132 did not affect the decrease in N protein expression induced by Neu5Ac2en-OAcOMe. Neu5Ac2en-OAcOMe also did not alter the levels of ubiquitination on HCoV-OC43-N protein (Figure. S3b). Thus, decreased N protein expression was not likely due to protein degradation.

### Antiviral activity of Neu5Ac2en-OAcOMe

Interestingly, Neu1 overexpression reversed the inhibition of Neu5Ac2en-OAcOMe on HCoV-OC43 infection (Fig. 4h), indicating Neu1 is the target of Neu5Ac2en-OAcOMe. Since Neu1 exists on the plasma membrane and cytoplasm(Chen et al., 2014a), we investigated whether Neu5Ac2en-OAcOMe-sensitive Neu1 resides on the plasma membrane or cytoplasm. To test this, we incubated HEK293T cells transfected with Neu1 expression vector with Neu5Ac2en-OAcOMe at room temperature for 30 min. We then measured cell surface sialidase activity and total cell lysate sialidase activity. Neu5Ac2en-OAcOMe significantly decreased total sialidase activity but did not affect cell surface sialidase activity (Fig. 4i), indicating that Neu5Ac2en-OAcOMe-sensitive Neu1 was located in the cytoplasm, where coronavirus replication takes place(Knoops et al., 2008). By contrast, Neu5Gc2en targets sialidase on cell surface(Chen *et al*., 2014a), which may explain why Neu5Gc2en did not inhibit HCoV-OC43 replication in THP-1 cells (Fig. 3a) or protect against SARS-CoV-2 infection-induced cell death (Fig. 3c). Consistent with previous research(Hata *et al*., 2008), oseltamivir treatment only slightly decreased sialidase activity in the cytoplasm (Fig 4j), as it is a prodrug as an ethyl ester.

Next we sought to ensure that the observed efficacy of Neu5Ac2en-OAcOMe was not restricted to THP-1 cells. We evaluated the efficacy of Neu5Ac2en-OAcOMe in three epithelial cell lines (HEK293T cells, HeLa cells and BSC-1 cells). Neu5Ac2en-OAcOMe inhibited viral replication in all three cell lines (Fig. 5a) and significantly decreased N protein levels (Fig. 5b). Immunofluorescence staining for HCoV-OC43-N protein also showed Neu5Ac2en-OAcOMe treatment effectively suppressed viral replication in BSC-1 cells (Fig. 5c). These results indicate the tested sialidase inhibitors did not inhibit the entry step of viral replication but interfered in the subsequent steps of the viral life cycle of coronavirus. Therefore, sialidase inhibitors represent a potential treatment for coronavirus infections.

**Figure 5.**
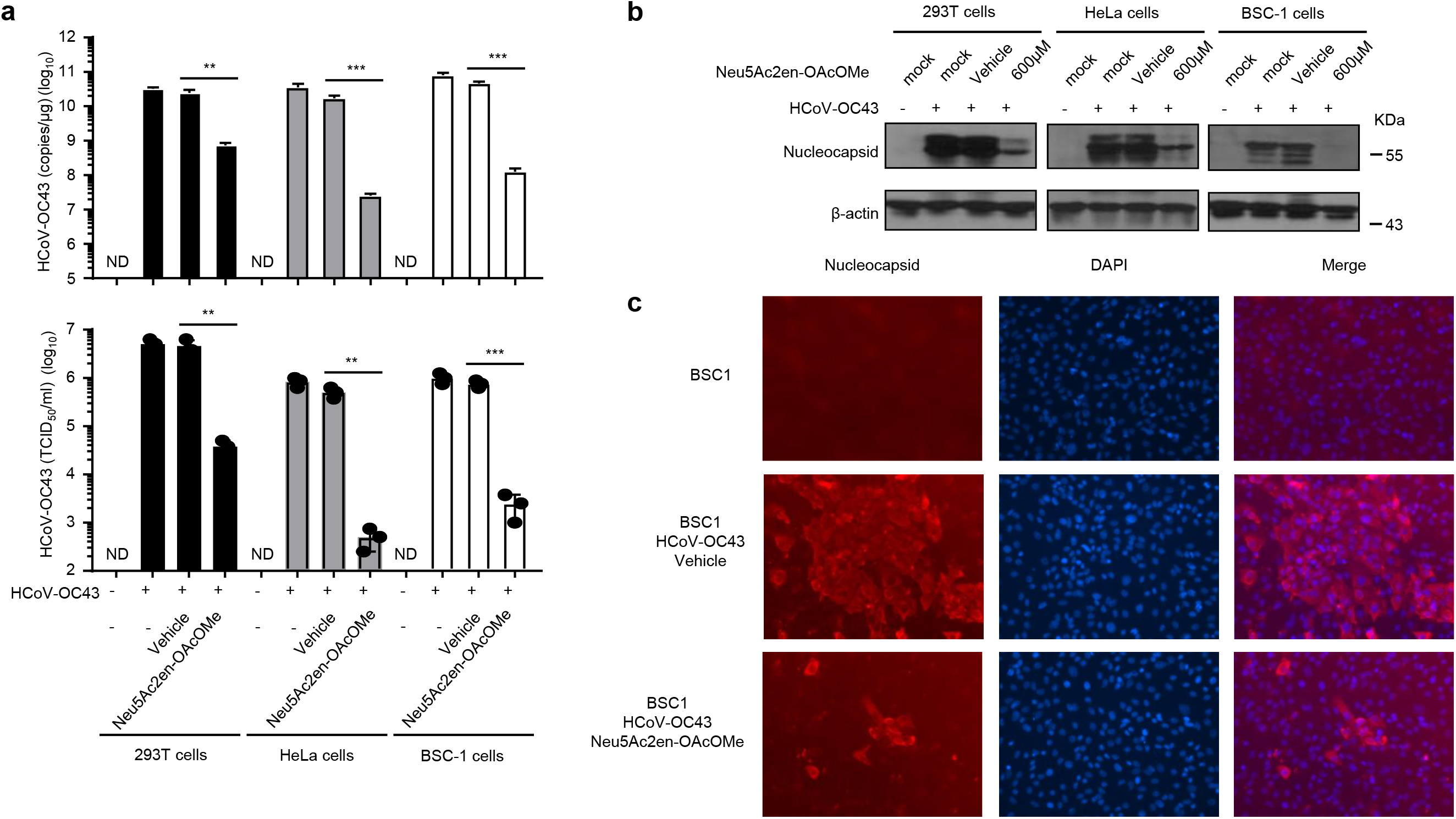
Neu5Ac2en-OAcOMe exhibits robust antiviral activity for coronavirus in epithelial cell lines. **a**, **b**, HEK293T (MOI = 1), HeLa (MOI = 1) and BSC-1 (MOI = 0.1) cells were inoculated with HCoV-OC43 virus for 2 hours, and Neu5Ac2en-OAcOMe was added after removal of inoculum. The levels of intracellular viral RNA (**a**, upper), virus release (**a**, lower), and intracellular nucleocapsid (**b**) were analyzed by RT-qPCR (**a**, upper), TCID_50_ assay (**a**, lower), and immunoblot (**b**), respectively (72 hours post-infection, except BSC-1, 48 hours post-infection). **c**, BSC-1 (MOI = 0.1) cells were inoculated with HCoV-OC43 virus for 2 hours, and Neu5Ac2en-OAcOMe was added after removal of inoculum. The levels of intracellular nucleocapsid were analyzed by immunofluorescence (48 hours post-infection). Red indicates nucleocapsid and blue represents cell nuclei stained by DAPI. Data are representative of three (**a, c**) or two (**b**) independent experiments. **p < 0.01; ***p < 0.001. ND, not detected. Analysis was performed using unpaired Student’s t-test.

### Therapeutic activity of Neu5Ac2en-OAcOMe

We further examined the antiviral efficacy of Neu5Ac2en-OAcOMe in vivo. No signs or symptoms of toxicity were observed during the treatment period (Figure. S4), so we determined whether Neu5Ac2en-OAcOMe could prevent HCoV-OC43 infection-induced death in newborn mice. Seven-day-old C57BL/6 pups were injected (intraperitoneal, IP) with 30 μl virus dilution (1 × 10^5^ TCID_50_ of HCoV-OC43). Based on our in vitro data, we chose to inject (IP) Neu5Ac2en-OAcOMe (20 mg/kg) 1 hour before virus infection. As shown in Fig. 6a, 100% of vehicle-treated mice succumbed to HCoV-OC43 challenge, while 50% of Neu5Ac2en-OAcOMe-treated mice survived throughout the observation period. Body weight was significantly lower in vehicle-treated mice than in Neu5Ac2en-OAcOMe-treated mice (Fig. 6b). Neu5Ac2en-OAcOMe-treated mice also showed suppression of HCoV-OC43 viral replication in the lungs, blood and brain (Fig. 6c).

**Figure 6.**
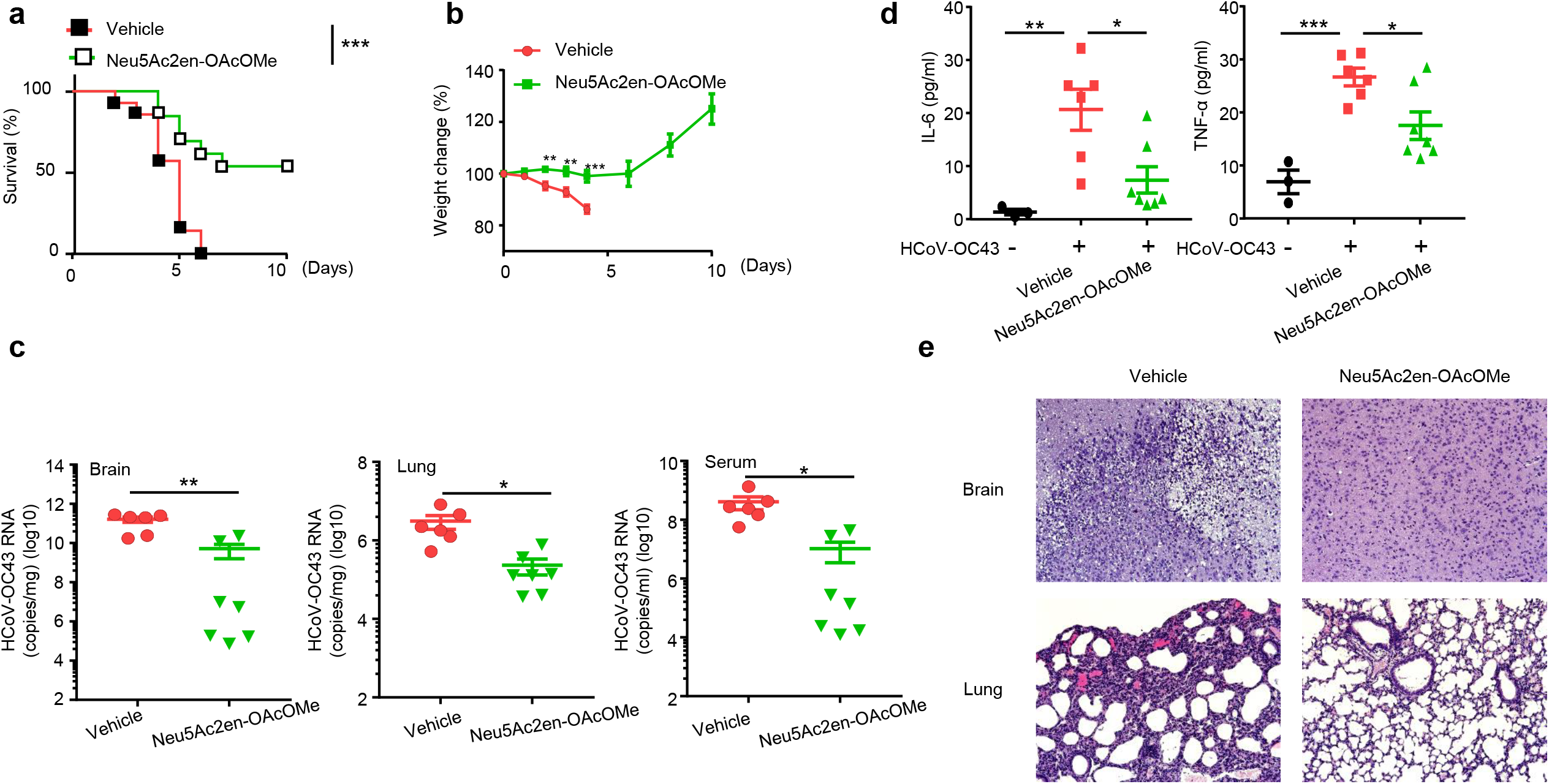
Evaluation of the therapeutic effects of Neu5Ac2en-OAcOMe. **a**-**e**, Seven-day-old mice were pre-treated with Neu5Ac2en-OAcOMe or vehicle and then IP infected with 30 μl virus dilution (1 × 10^5^ TCID_50_ HCoV-OC43). **a**, Survival curves after HCoV-OC43 infection. (n=11 for vehicle group, n=10 for Neu5Ac2en-OAcOMe group). **b**, Body weight after HCoV-OC43 infection. (n=11 for vehicle group, n=10 for Neu5Ac2en-OAcOMe group). **c**, Viral RNA copies of HCoV-OC43 in the blood, brain and lung day 5 post-infection. (n=6 for vehicle group, n=7 for Neu5Ac2en-OAcOMe group). **d**, Cytokines in blood day 5 post-infection. (n=6 for vehicle group, n=7 for Neu5Ac2en-OAcOMe group). **e**, Histological analysis of brain and lung tissues day 5 post-infection. Tissue sections were stained with H&E. Data are representative of at least three independent experiments (**a**-**e**). *p < 0.05; **p < 0.01; ***p < 0.001. Analysis was performed using Kaplan Meier analysis (**a**), unpaired Student’s t-test (**b, c**), or one-way ANOVA (**d**).

Both viral and host factors impact disease pathogenesis. During infection with SARS-CoV, Middle East respiratory syndrome coronavirus and SARS-CoV-2, cytokine storm is a major cause of mortality(Channappanavar and Perlman, 2017; Moore and June, 2020). In our previous study, sialidase inhibitors rescued mice from bacterial infection-induced death by inhibiting the activity of host cell surface Neu1 and suppressing the cytokine storm(Chen *et al*., 2014a). To determine whether Neu5Ac2en-OAcOMe exerts similar effects in coronavirus infection, we measured serum levels of interleukin 6 (IL-6) and tumor necrosis factor alpha (TNF-α), which contribute to the cytokine storm and correlate with respiratory failure and adverse clinical outcome in COVID-19(Karki et al., 2021). We detected substantially decreased serum IL-6 and TNF-α levels in Neu5Ac2en-OAcOMe-treated mice (Fig. 6d). Hematoxylin and eosin (H&E) staining also indicated less brain and lung tissue damage in Neu5Ac2en-OAcOMe-treated mice (Fig. 6e). Taken together, these findings show Neu5Ac2en-OAcOMe conferred significant protection against HCoV-OC43 challenge by reducing viral replication in vivo and the associated inflammatory dysregulation.

## Discussion

N protein has a mass of 50 to 60 kDa (Fig. 1 and 2), indicating the presence of post-translational modifications such as N or O-linked glycosylation (Figure. S1) and sialylation (Fig. 1). Moreover, SARS-CoV-2-N protein is highly glycosylated, as demonstrated by glycomic and glycoproteomic analyses after expression in HEK293T cells(Supekar *et al*., 2021). In this study, N protein from SARS-CoV-2 and HCoV-OC43 was significantly sialylated and this sialylation was tightly regulated by host Neu1. Coronavirus replication occurs in the cytoplasm of infected host cells(Knoops *et al*., 2008). The sialidase inhibitor Neu5Ac2en-OAcOMe targets cytoplasmic sialidase but not cell surface sialidase, which is advantageous because β-Coronaviruses traffic to lysosomes(Blaess et al., 2020; Ghosh *et al*., 2020; Zheng et al., 2018) where Neu1 is also known to be predominantly localized(Miyagi and Yamaguchi, 2012). Notably, the newly developed sialidase inhibitor Neu5Ac2en-OAcOMe reduced HCoV-OC43 replication in vitro and in vivo by inhibiting host Neu1 activity and rescued mice from HCoV-OC43 infection induced death. Moreover, Neu5Ac2en-OAcOMe reduced SARS-CoV-2 replication in vitro.

Inhibition of viral neuraminidase activity has developed as a therapeutic approach for influenza infection(Glanz *et al*., 2018). Tamiflu (oseltamivir) and Relenza (Zanamivir), which are approved for treatment of influenza A and B, have almost no effect on human neuraminidases(Glanz *et al*., 2018; Hata *et al*., 2008). Several clinical trials have assessed the efficacy of oseltamivir in treating SARS-CoV-2 infection, but no positive outcomes were observed(Apaydin et al., 2021; Sanders et al., 2020; Tan et al., 2021; Yousefi et al., 2020; Zhao et al., 2021). This lack of efficacy could be attributed to several reasons: 1) SARS-CoV-2 genomic RNA does not encode sialidase(Mariano et al., 2020; Vandelli et al., 2020); 2) SARS-CoV-2 replication depends on the host Neu1 (Fig. 2); and 3) oseltamivir has few inhibitory effects on host Neu1 (Fig. 4j)(Hata *et al*., 2008). Together, these findings explain why oseltamivir has not shown efficacy in the treatment of COVID-19. Here, we demonstrate that a newly synthesized sialidase inhibitor Neu5Ac2en-OAcOMe which specifically targets intracellular host sialidase Neu1 and inhibits coronavirus replication (Fig. 7), accounting for the contributions of both the host and pathogen in the disease process. Based on our findings, sialidase inhibitors could be a generalizable and effective treatment in the current COVID-19 pandemic as well as future coronavirus pandemics associated with the inflammatory response.

**Figure 7.**
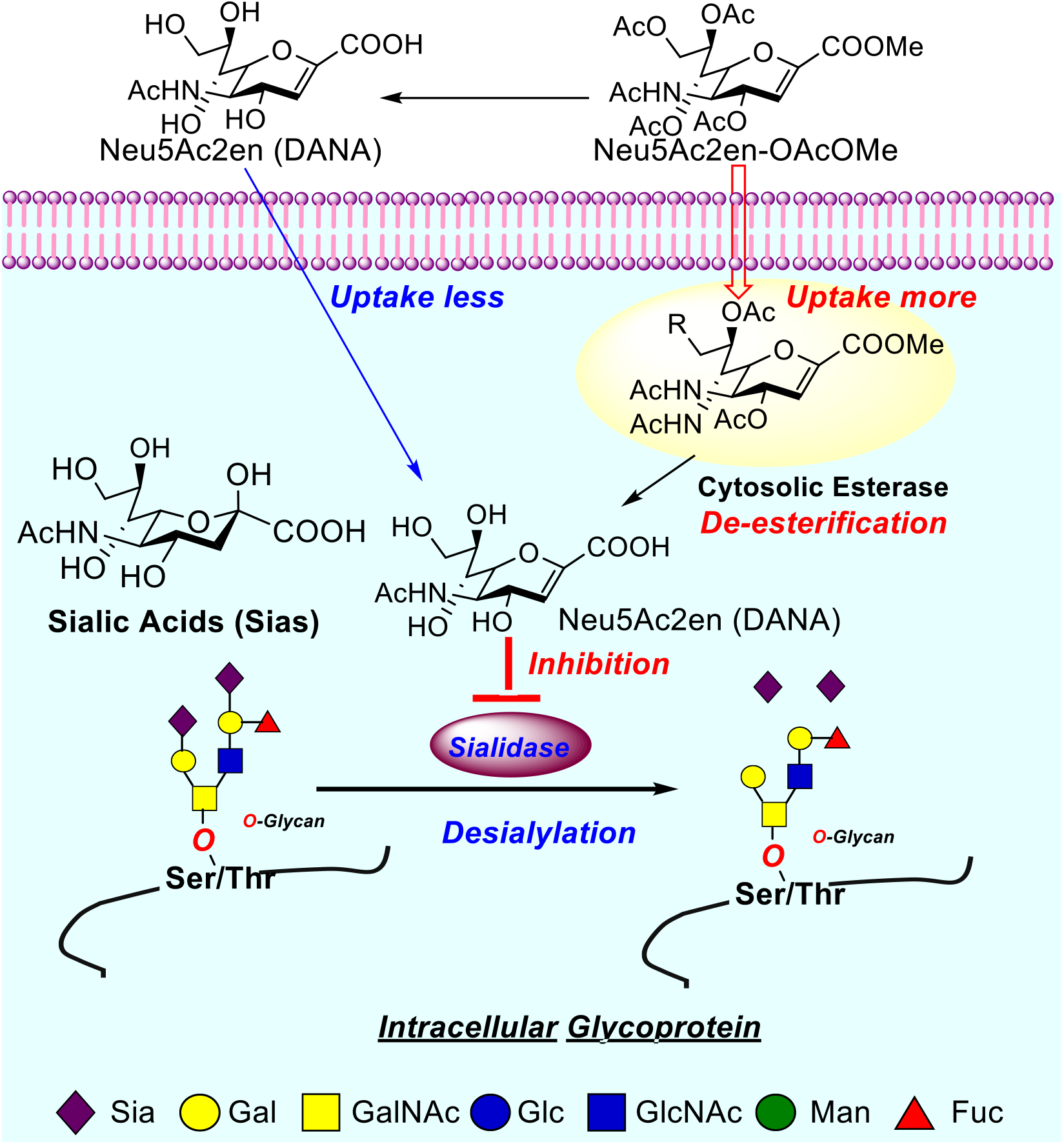
Proposed inhibition of cytosolic desialylation (sialidase) by cytosolic sialidase inhibitor.

## Author contributions

D.Y., Y.W., I. T., J.K. and G.Y.C. performed experiments and analyzed and/or interpreted results. X.L.S supervised synthesis of sialidase inhibitors. K.L. provided study materials, expertise and feedback. M.C analyzed data and edited the text of a draft of this manuscript. R.L. and L.W. provided expertise and feedback. G.Y.C. designed and supervised the overall study. D.Y. and G.Y.C. wrote the paper.

## STAR Methods

### Reagents

COVID-19 patient sera (both IgG and IgM antibodies to the N protein were negative) were purchased from Raybiotech (Peachtree, GA). Anti-SARS-CoV-2 N protein (HL5410, Catalog # MA5-36270) was obtained from Thermo Fisher Scientific (Waltham, MA). Anti-HCoV-OC43 N protein and anti-human Neu1 anti-antibodies were purchased from Sigma-Aldrich (St. Louis, MO). Anti-ubiquitin mouse monoclonal antibody (FK2) (cat. no. ST1200, lot no. D00165221) was obtained from EMD Millipore (Merck KGaA, Darmstadt, Germany). MG132 (cat. no. 3175-v, lot no. 640311) was purchased from Peptide institute (Osaka, Japan). Biotinylated Maackia Amurensis Lectin II (MAL II, MAA) (cat. no. B-1265) and Biotinylated Sambucus Nigra Lectin (SNA, EBL) (cat. no. B-1305) were purchased from Vector Laboratories (Burlingame, CA). Anti-β-actin, Streptavidin-Horseradish peroxidase (HRP) and HRP-conjugated anti-mouse, anti-goat or anti-rabbit secondary antibodies were purchased from Santa Cruz Biotechnology (Santa Cruz, CA). Human Neu1 shRNAs were purchased from Sigma. HeLa, BSC-1, HEK293T and THP-1 cells were obtained from ATCC (Manassas, VA) and cultured in Dulbecco’s Modified Eagle Medium (DMEM) or Roswell Park Memorial Institute (RPMI) medium supplemented with 10% heat-inactivated fetal bovine serum, 2 mM glutamine, and 100 μg/ml penicillin/streptomycin. Neuraminidase (sialidase) provided by Vibrio cholerae (cat. no. 11080725001) was purchased from Sigma. The 32m ssDNA (5’-CGAGGCCACGCGGAGTACGATCGAGGGTACAG-3’) was purchased from Thermo Fisher Scientific. The 32m ssRNA (5’-CGAGGCCACGCGGAGUACGAUCGAGGGUACAG-3’) was purchased from Eurofins Genomics (Louisville, KY). SARS-CoV-2 nucleocapsid encoding plasmid was purchased from Sino Biological (cat. No. VG40588-UT, Beijing, China). Oseltamivir, Zanamivir and Neu5Gc2en were obtained from Thermo Fisher Scientific. Neu5Ac2en9N3, Neu5Ac2en-OMe, Neu5Ac2en-OAcOMe and Neu5Ac2en9N3-OAcOMe were synthesized as described(Sun et al., 2000; Zou et al., 2010).

### Syntheses of Sialidase Inhibitors

Neu5Ac2en-OMe and Neu5Ac2en-OAcOMe were synthesized using our previously reported method(Sun *et al*., 2000).

Neu5Ac2en9N3 is synthesized by literature method(Zou *et al*., 2010).

Neu5Ac2en9N3-OAcOMe is newly made from Neu5Ac2en9N3-OMe(Zou *et al*., 2010).

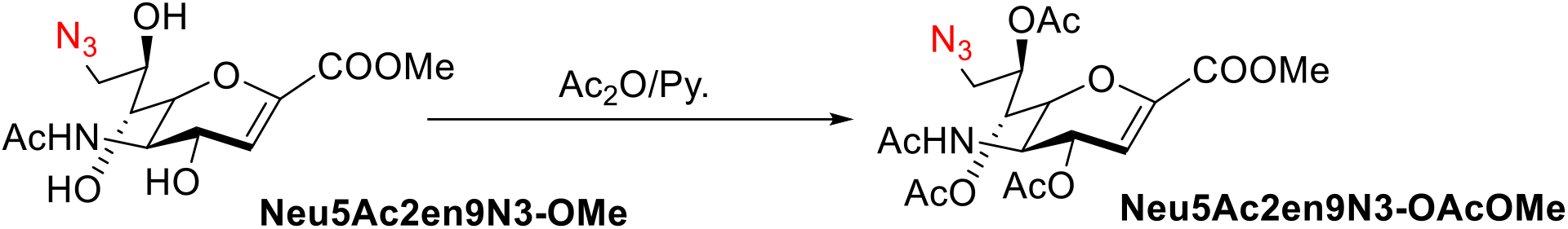

Neu5Ac2en9N3-OMe (100 mg) was dissolved into a minimal amount of a 1:3 solution of acetic anhydride in anhydrous pyridine at 0 °C, then warmed to room temperature and stirred for 18 hours. The solution was concentrated and the residue was purified by flash chromatography with 5:1 EtOAc-hexane as the mobile phase to give Neu5Ac2en9N3-OAc-OMe as a white solid (65 % yield,). ^1^H NMR (400 MHz, CD_3_OD): δ 6.05 ppm (d, J=3 Hz, 1H), 5.53 ppm (dd, J= 3 Hz, 1H), 5.51 ppm (dd, J= 3 Hz, 1H), 5.23 ppm (m, 1H), 4.39 ppm (dd, J= 9, 3 Hz, 2H), 3.95 ppm (dd, J=12, 2 Hz, 1 H), 3.86 ppm (s, 3H), 3.54 ppm (dd, J=8, 3 Hz, 1 H), 2.12 ppm (s, 3H), 2.08 ppm (s, 3H), 2.05 ppm (s, 3H), 1.89 ppm (s,3H). ^13^C NMR (101 MHz, CDCl3): δ 172.3, 171.5, 162.3, 147.3, 108.6, 73.4, 67.8, 52.3, 50.5, 46.9, 23.5, 21.8 (Figure. S5).

### Construction of plasmids

To generate the constructs expressing human Neu1, cDNA for Neu1 was amplified by RT-PCR and subcloned into expression vectors pcDNA6 and pLVX-puro (Life Technologies, Carlsbad, CA). All constructs were verified by restriction enzyme digestion and DNA sequencing.

### Gel-mobility-shift assay

ssDNA or ssRNA in phosphate buffer (10 mM sodium phosphate, 50 mM NaCl, 1 mM EDTA, 0.01% NaN3, pH 7.4) was heated to 95°C and immediately put on ice to destroy its secondary structure. 1x 10^7^ HEK293T cells in a 10 cm dish transfected with empty vector or SARS-CoV-2 N protein expression vector for 48 hours were harvested, suspended in lysis buffer (20 mM Tris-HCl, 0.1 % Triton X-100, 150 mM NaCl, pH 7.6) and separated equally. Half of the cell lysates were treated with sialidase for 2 hours at 37°C. The oligonucleotides were mixed with the cell lysates and incubated on ice for 10 min and then separated on 1% agarose gels. For antibody supershift assays, 1μg of anti-N protein antibodies was added to the reaction mixture and then separated in a 7% acrylamide gel as described in previous publication (Chen et al., 2004).

### Experimental animal models

WT C57BL/6J mice were obtained from The Jackson Laboratory (Bar Harbor, ME). All animal procedures were approved by the Animal Care and Use Committee of University of Tennessee Health Science Center. The HCoV-OC43 infection mouse model was established as described previously(Jacomy and Talbot, 2003). Briefly, 7-day-old mice were separated randomly into two groups and injected intraperitoneally (IP) with either Neu5Ac2en-OAcOMe (20 mg/kg) or vehicle (0.5% dimethyl sulfoxide, DMSO). One hour later, mice were inoculated with 30 μl of virus dilution (1 x 10^5^ TCID_50_ of HCoV-OC43) by IP injection. Neu5Ac2en-OAcOMe and vehicle were administered daily and mice were monitored up to 10 days for survival. To detect viral RNA loads in tissues and cytokine production, mice were euthanized at 5 days post-infection. Mouse brain, lung, and blood tissues were collected.

### Immunofluorescence

Cells were fixed with 4% paraformaldehyde in phosphate buffered saline (PBS) at room temperature for 15 min and then permeabilized with 1% Triton X-100 in PBS at room temperature for 15 min. Immunofluorescence staining was performed as described previously(Yang et al., 2019). Images were acquired with an EVOS FL Auto Imaging System (Thermo Fisher Scientific).

### HCoV-OC43 production and titration

HCoV-OC43 virus (ATCC^®^ VR-1558™) was purchased from ATCC. The stock of HCoV-OC43 was produced and titrated using BSC-1 cells. Viral titers in cell-free culture supernatants were determined by endpoint dilution-based TCID_50_ assays in 96-well plates(Liu *et al*., 2014). Cytopathic effect was recorded and used for calculation of viral titers at 7 days post-infection.

### SARS-CoV-2 high-throughput screen (HTS) cytopathic effect assay

Double-blinded SARS-CoV-2 high-throughput screen (HTS) cytopathic effect assay was performed with the fee service provided by the University of Tennessee Health Science Center Regional Biocontainment BSL3 Laboratory. The basic methods for the HTS for the identification of potential inhibitors of coronavirus have been previously described(Severson et al., 2007). Briefly, seven sialidase inhibitors (oseltamivir, Zanamivir, Neu5Gc2en, Neu5Ac2en9N3, Neu5Ac2en-OMe, Neu5Ac2en-OAcOMe and Neu5Ac2en9N3-OAcOMe) were plated in 384-well black wall plates containing 4,500 Vero 76 cells/well in single dose of indicated concentration in Eagle’s MEM with 5% heat inactivated FBS, 1% penicillin/streptomycin/L-glutamine, 1% HEPES and 0.5% DMSO. The cells were infected with SARS-CoV-2 at an MOI of 0.1. Plates were then incubated at 37°C, 5% CO2, for 48 hours. The cell viability at the end of incubation period was measured as described elsewhere(Chung et al., 2010). After incubation, 100 μL of Promega CellTiter-GloR (Promega, Madison, WI) was added to each well using the BiomekR 2000. Plates were shaken for 2 min at speed 5 on a Labline Instruments (Kochi, India) plate shaker. Luminescence was then measured using a PerkinElmer Envision™ plate reader (PerkinElmer, Wellesley, MA).

### Neuraminidase activity assay

Sialidase activity was measured using 2’-(4-methylumbelliferyl)-α-D-N-acetylneuraminic acid sodium salt hydrate (4-MU-NANA, catalog no. sc-222055, Santa Cruz Biotechnology) as the substrate. 1 x 10^7^ HEK293T cells in a 10 cm dish transfected with Neu1 expression vector were harvested after 48 hours, incubated with inhibitors for 30 min at room temperature, washed to remove the sialidase inhibitors, and separated equally. Half of the cells were used for detection of cell surface sialidase activity and half of the cells were suspended in lysis buffer (20 mM Tris-HCl, 0.1 % Triton X-100, 150 mM NaCl, pH 7.6) for detection of whole cell sialidase activity. For the reaction, intact cells or lysed cells were incubated with 4-MU-NANA (final concentration, 15 μM) for 30 min at 37°C in 50 μl reaction buffer (50 mM Sodium phosphate, pH 5.0). The reaction was terminated by adding 600 μl stop buffer (0.25 M glycine-NaOH, pH 10.4). Fluorescence intensity was measured with a Synergy HTX Multi-Mode Reader (EMD Millipore, Merck KGaA) (excitation 360 nm; emission 460 nm).

### Real-time quantitative PCR

Total RNA was extracted with TRIzol (Invitrogen, Carlsbad, CA) according to the manufacturer’s protocol and reverse transcribed with random primers and Superscript III (Life Technologies). The mRNA expression of human Neu1, Neu2, Neu3 and Neu4 was measured by real-time PCR. Samples were run in triplicate, and the relative expression was determined by normalizing the expression of each target to the endogenous reference, GAPDH. For RT-qPCR, copy numbers of HCoV-OC43 viral RNA were calculated based on a standard curve generated using pEF6-OC43-N-V5 His DNA. The following primers were used: Neu1: 5’-GGAGGCTGTAGGGTTTGGG-3’ (forward), 5’-CACCAGACCGAAGTCGTTCT-3’ (reverse); Neu2: 5’-CCATGCCTACAGAATCCCTGC-3’ (forward), 5’-CTCTGCGTGCTCATCCTTC-3’ (reverse); Neu3: 5’-AAGTGACAACATGCTCCTTCAA-3’ (forward), 5’-TCTCCTCGTAGAACGCTTCTC-3’ (reverse); Neu4: 5’-GGCCACGGGATGACAGTTG-3’ (forward), 5’-CAGGCGGATACCCATGTGAG-3’ (reverse); HCoV-OC43 N gene, 5’-CGATGAGGCTATTCCGACTAGGT-3’ (forward) and 5’-CCTTCCTGAGCCTTCAATATAGTAACC-3’ (reverse).

### Immunoblotting

Cell lysates were prepared in lysis buffer (20 mM Tris-HCl, 150 mM NaCl, 1 % Triton X-100, pH 7.6, including protease inhibitors, 1 μg/ml leupeptin, 1 μg/ml aprotinin and 1 mM phenylmethylsulfonyl fluoride), sonicated, centrifuged at 13,000 rpm for 5 min and then analyzed by Western blot. The concentration of running gel was 10%. After blocking, the blots were incubated with the appropriate primary antibody (1: 1,000 dilution) or Biotin-MAA/SNA (1 μg/ml). After incubation with the secondary antibody (HRP-conjugated goat anti-rabbit IgG, goat anti-mouse IgG, 1: 5,000 dilution) or Streptavidin-HRP (1: 10,000 dilution), the signal was detected with an enhanced chemiluminescence (ECL) kit (Santa Cruz).

### Measurement of inflammatory cytokines

Mouse blood samples were obtained at indicated time points, and cytokines in the serum were determined using a mouse cytokine bead array designed for inflammatory cytokines (552364, BD Biosciences, San Jose, CA). Human cytokines in cell culture-derived supernatants were determined using a human cytokine bead array designed for inflammatory cytokines (551811, BD Biosciences).

### Statistical analysis

GraphPad Prism software (San Jose, CA) was used for data analysis. Data are shown as mean ± SD or mean ± SEM. Statistical significance was analyzed by two-tailed t-test for two groups or one-way analysis of variance (ANOVA) or two-way ANOVA for three or more groups. Differences in survival rates were analyzed by Kaplan-Meier plot and statistical significance was determined using a log-rank (Mantel-Cox) test. *P<0.05, **P<0.01, ***P<0.001, n.s., not significant.

