## Supplemental data for "Targeting intracellular Neu1 for Coronavirus Infection Treatment"

**Figure. S1. HCoV-OC43 nucleocapsid is an N-linked (a) and O-linked (b) glycosylation protein.**

HCoV-OC43 nucleocapsid was isolated and purified from HCoV-OC43 virion by SDS-PAGE. The band for nucleocapsid was cut out from SDS-PAGE gel, confirmed by Western blotting and then sent to the Harvard Glycomics Core for N- and O-Glycan profiling.

**Figure. S2. Neu5Ac2en-OAcOMe treatment did not induce TNF-α, IL-6 and IL-1β production in THP-1 cells.**

THP-1 cells with or without HCoV-OC43 (MOI = 2) infection for 72 hours or treated with poly(IC) (100 μ/ml) or LPS (0.2 μ/ml) for 18 hours. Cytokines in the cell culture supernatants were determined with a Human Inflammatory Cytokine kit. LPS and Poly(IC) were used as positive controls. ***$p < 0.001$. Analysis was performed using one-way ANOVA.

**Figure. S3. Neu5Ac2en-OAcOMe treatment did not affect the stability of N protein.**

**a**, THP-1 cells infected with HCoV-OC43 were treated with or without Neu5Ac2en-OAcOMe and MG132. Nucleocapsid in the cell lysates was measured by western blot. **b**, THP-1 cells infected with HCoV-OC43 and treated with or without Neu5Ac2en-OAcOMe were immunoprecipitated with an anti-N Ab and blotted for ubiquitin (FK2 Ab), and anti-N antibodies.

**Figure. S4. Toxicity test of Neu5Ac2en-OAcOMe.**

The mice treated with Neu5Ac2en-OAcOMe did not show any signs or symptoms of toxicity during the treatment period. There was no weight loss from Day 1 to Day 4 after treatment. There was no significant induction of TNF-α, IFN-γ or IL-6 during treatment. The mice were treated with the Neu5Ac2en-OAcOMe for 4 days survived 2 weeks, and all mice showed normal activity. Three 5-6 week-old females were IP injected with Neu5Ac2en-OAcOMe (20 mg/kg) daily for 4 days. Day 0 is untreated and body weight was set as 100%.

**Figure. S5. NMR spectra of Neu5Ac2en9N3-OAcOMe (400 MHz, $CDCl_3$).**

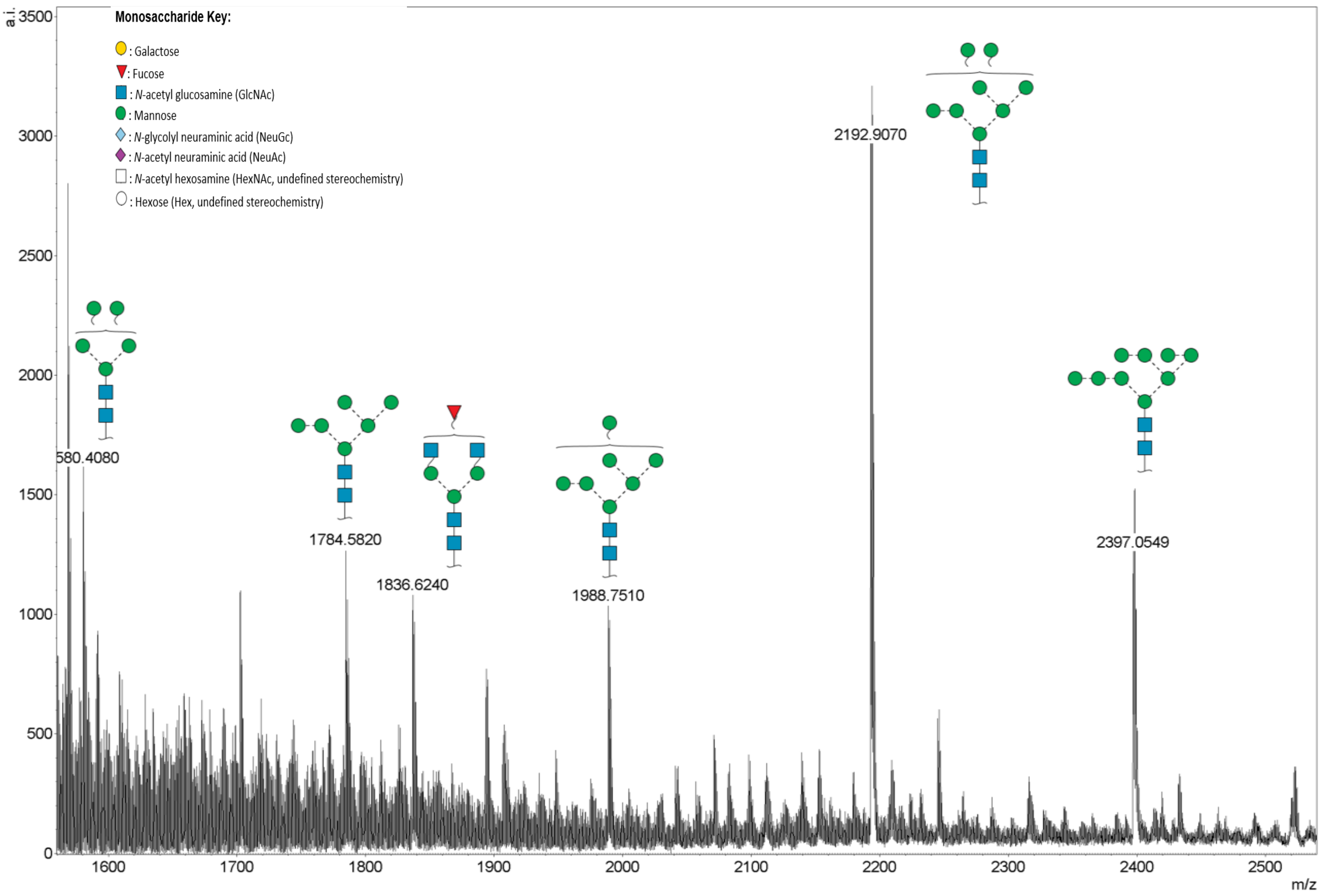

Monosaccharide Key:
: Galactose
: Fucose
: N-acetyl glucosamine (GlcNAc)
: Mannose
: N-glycolyl neuraminic acid (NeuGc)
: N-acetyl neuraminic acid (NeuAc)
: N-acetyl hexosamine (HexNAc, undefined stereochemistry)
: Hexose (Hex, undefined stereochemistry)
a.i.
3500
3000
2500
2000
1500
1000
500
0
580.4080
1784.5820
1836.6240
1988.7510
2192.9070
2397.0549
1600
1700
1800
1900
2000
2100
2200
2300
2400
2500
m/z


Figure. S1a

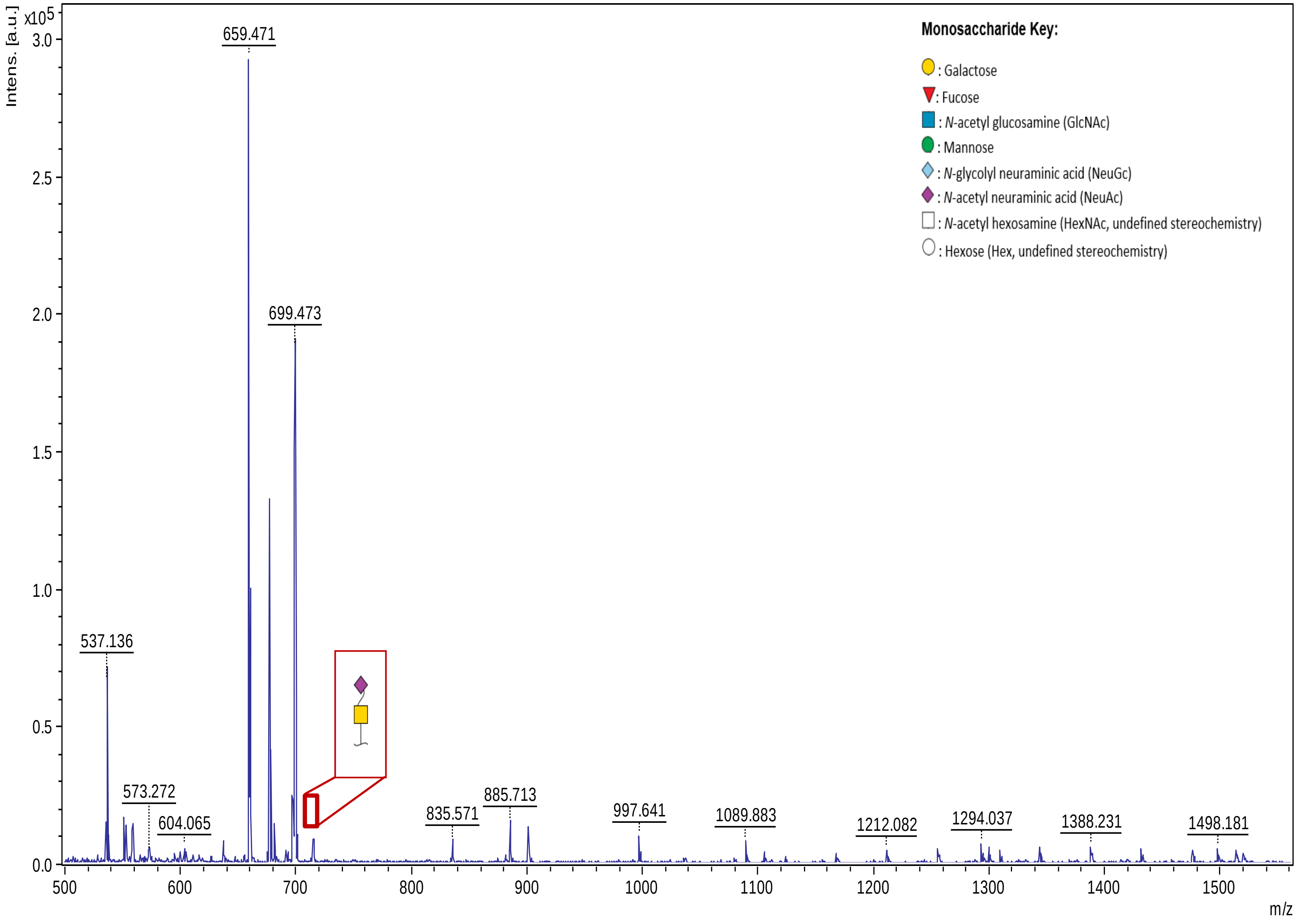

Intens. [a.u.]
x10^5
3.0
2.5
2.0
1.5
1.0
0.5
0.0
659.471
699.473
537.136
573.272
604.065
835.571
885.713
997.641
1089.883
1212.082
1294.037
1388.231
1498.181
500
600
700
800
900
1000
1100
1200
1300
1400
1500
m/z
Monosaccharide Key:
: Galactose
: Fucose
: N-acetyl glucosamine (GlcNAc)
: Mannose
: N-glycolyl neuraminic acid (NeuGc)
: N-acetyl neuraminic acid (NeuAc)
: N-acetyl hexosamine (HexNAc, undefined stereochemistry)
: Hexose (Hex, undefined stereochemistry)


Figure. S1b

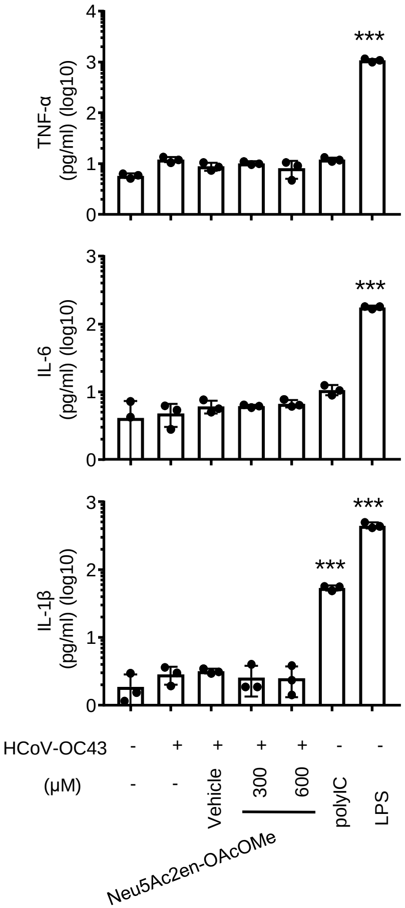
TNF-α
(pg/ml) (log10)
IL-6
(pg/ml) (log10)
IL-1β
(pg/ml) (log10)
***
HCoV-OC43
(μM)
Vehicle
300
600
polyIC
LPS
Neu5Ac2en-OAcOMe

Figure. S2

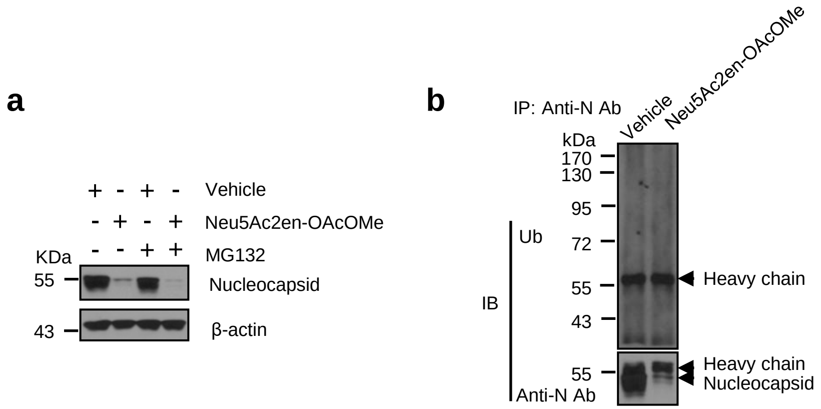
a
+ - + - Vehicle
- + - + Neu5Ac2en-OAcOMe
KDa - - + + MG132
55
Nucleocapsid
43
β-actin
b
IP: Anti-N Ab
Vehicle
Neu5Ac2en-OAcOMe
kDa
170
130
95
72
55
43
Ub
IB
Heavy chain
55
Heavy chain
Nucleocapsid
Anti-N Ab

Figure. S3

a

b

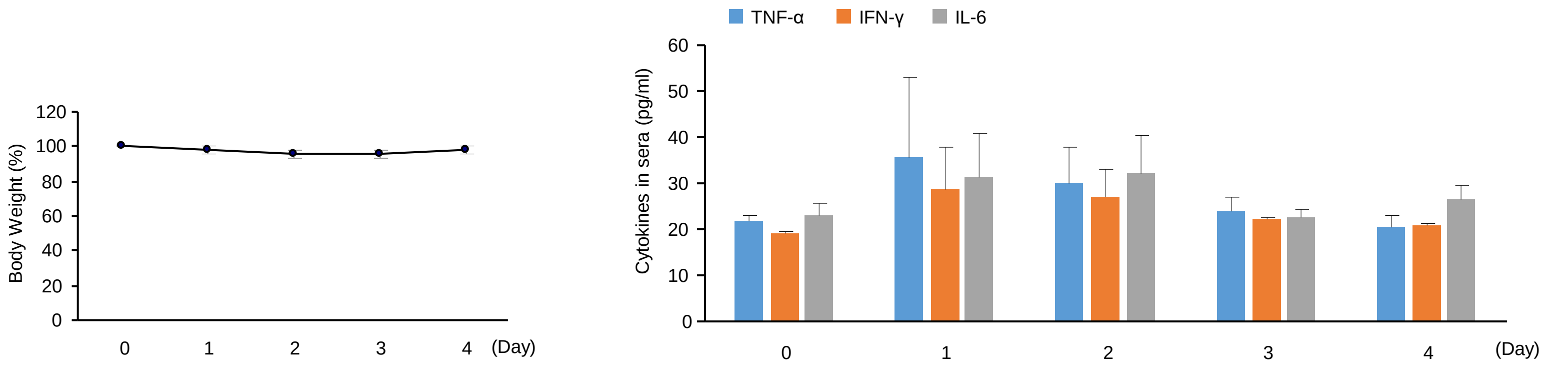
Body Weight (%)
120
100
80
60
40
20
0
0
1
2
3
4
(Day)
TNF-α
IFN-γ
IL-6
Cytokines in sera (pg/ml)
60
50
40
30
20
10
0
0
1
2
3
4
(Day)

Figure. S4

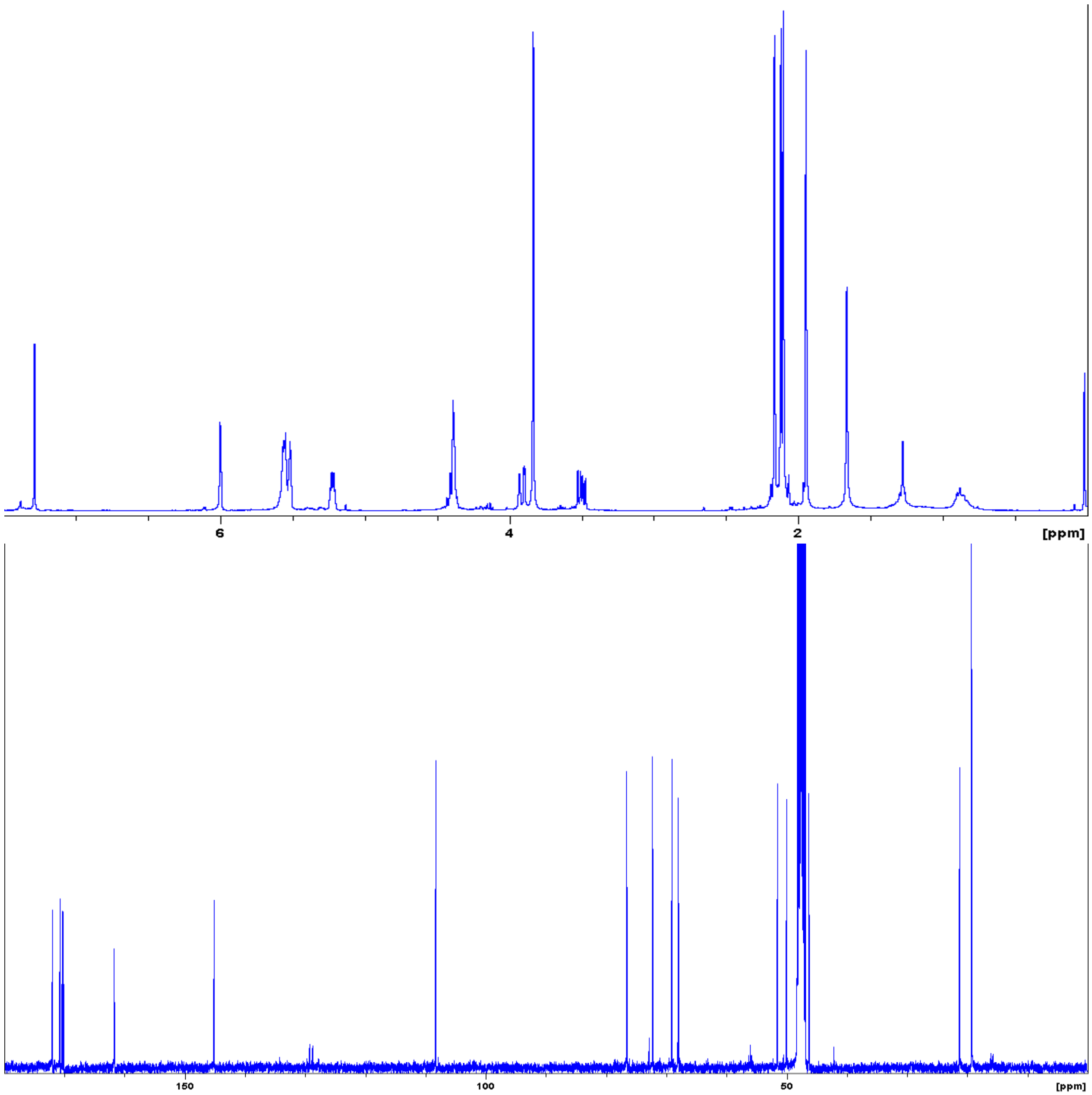


Figure. S5
